## Supplementary Material for "The specific features of the developing T cell compartment of the neonatal lung are a determinant of respiratory syncytial virus immunopathogenesis"

**Table S1. List of antibodies validated for their use in the study.**

| Antibody | Dilution | Host, isotype | Clone | Reference, Source |
|---|---|---|---|---|
| Anti-DEC205 | 1:20 | Mouse, IgG2b | CC98 | MCA1651GA, BioRad |
| Anti-CD11c | 1:33 | Mouse, IgM | BAQ153A | WS0519B-100, Kingfisher |
| Anti-CD172a | 1:400 | Mouse, IgG1 | DH59B | WS0567B-100, Kingfisher |
| Anti-CD13 | 1:100 | Mouse, IgG1 | CC81 | MCA2338GA, BioRad |
| Anti-CD86-PE | 1:10 | Mouse, IgG1 | IL-A190 | MCA2437PE, Bio-Rad |
| Anti-IL-12/23 | 1:80 | Mouse, IgG2a | CC301 | MCA1782EL, Bio-Rad |
| Anti-CD45RO-PE | 1:10 | Mouse, IgG3 | IL-A116A | MCA2434PE, Bio-Rad |
| Anti-CD4 | 1:400 | Mouse, IgG2a | 44.38 | MCA2213GA, Bio-Rad |
| Anti-CD8 | 1:160 | Mouse, IgG1 | CC58 | MCA1654G, Bio-Rad |
| Anti-CD25 | 1:200 | Mouse, IgG3 | LCTB2A | WS0597B-100, Kingfisher |
| Anti-IFN-γ-AF647 | 1:160 | Mouse, IgG1 | CC302 | MCA1783A647, Bio-Rad |
| Anti-IL-4-FITC | 1:10 | Mouse, IgG2a | CC303 | MCA1820F, Bio-Rad |
| Anti-FOXP3-FITC | 1:50 | Rat, IgG2a | FJK-16s | 11-5773-82, TermoFisher |
| Anti-TGF-β | 1:100 | Mouse, IgG1 | TB21 | MCA797, Bio-Rad |
| Anti-IL-10 | 1:40 | Mouse, IgG2b | CC318 | MCA2110, Bio-Rad |
| Anti-WC1-FITC | 1:80 | Mouse, IgG2a | CC15 | MCA838F, Bio-Rad |
| Anti-IL-17A-PE | 1:20 | Mouse, IgG1 | Ebio64DEC17 | 12-7179-42, ThermoFisher |
| Anti-mouse IgG1, FITC | 1:1000 | Goat, IgG | - | A-21121, ThermoFisher |
| Anti-mouse IgG1, PE | 1:200 | Goat, IgG | - | P-21129, ThermoFisher |
| Anti-mouse IgG1, PerCP-Cy5.5 | 1:80 | Rat, IgG | RMG1-1 | 406612, Biolegend |
| Anti-Mouse IgG3, PerCP-Cy5.5 | 1:200 | Goat, IgG | - | 1100-13, SouthernBiotech |
| Anti-mouse IgM, PerCP-Cy5.5 | 1:20 | Rat, IgG2a | RMM-1 | 406512, Biolegend |
| Anti-mouse IgG1, PE-Cy7 | 1:640 | Rat, IgG | RMG1-1 | 406614, Biolegend |
| Anti-mouse IgG2b, PE-Cy7 | 1:250 | Goat, IgG | - | 1090-17, SouthernBiotech |
| Anti-mouse IgG1, a647 | 1:1000 | Goat, IgG | - | A-21240, ThermoFisher |
| Anti-mouse IgG2b, a647 | 1:1000 | Goat, IgG | - | A-21242, ThermoFisher |
| Anti-mouse IgG1, APC-Cy7 | 1:40 | Rat, IgG | RMG1-1 | 406620, BioLegend |
| Anti-mouse IgG2a, APC-Cy7 | 1:250 | Goat, IgG | - | 1080-19, SouthernBiotech |
| Anti-mouse IgG1, BV421 | 1:80 | Rat, IgG | RMG1-1 | 406616, BioLegend |

### Table S2. Combination immunostainings used for the identification of cell subsets.

| Subset | Live/Dead Aqua | Surface 1 | Surface 2 | IgG block | Surface 3 | Fix & Perm | Intracell. 1 | Intracell. 2 |
|---|---|---|---|---|---|---|---|---|
| **pDC** | + | CD172a-IgG1<br>CD205-IgG2b<br>CD11c-IgM | IgG1-PE-Cy7<br>IgG2b-AF647<br>IgM-PerCP-Cy5 | + | CD13-IgG1-AF488<br>CD86-IgG1-PE | + | TNF-IgG2a | IgG2a-APC-Cy7 |
| **Th2/Tc2** | + | CD8-IgG1<br>CD4-IgG2a<br>CD25-IgG3 | IgG1-PE-Cy7<br>IgG2a-APC-Cy7<br>IgG3-PerCP-Cy5.5 | + | CD45RO-IgG3-PE | + | IL-4-IgG2a-FITC<br>IFN-γ-IgG1-AF647 | - |
| **γδ T** | + | CD8-IgG1<br>CD4-IgG2a<br>CD25-IgG3 | IgG1-PE-Cy7<br>IgG2a-APCy-Cy7<br>IgG3-PerCP-Cy5.5 | + | WG1-IgG1-FITC | + | IL17A-IgG1-PE<br>IFN-γ-IgG1-AF647 | - |
| **Tregs** | + | CD4-IgG2a<br>CD25-IgG3 | IgG2a-APC-Cy7<br>IgG3-PerCP-Cy5.5 | + | CD45RO-IgG3-PE | + | TGF-β-IgG1<br>IL10-IgG2b | IgG1-PE-Cy7<br>IgG2b-AF647<br>FOXP3-IgG2a-AF488 |

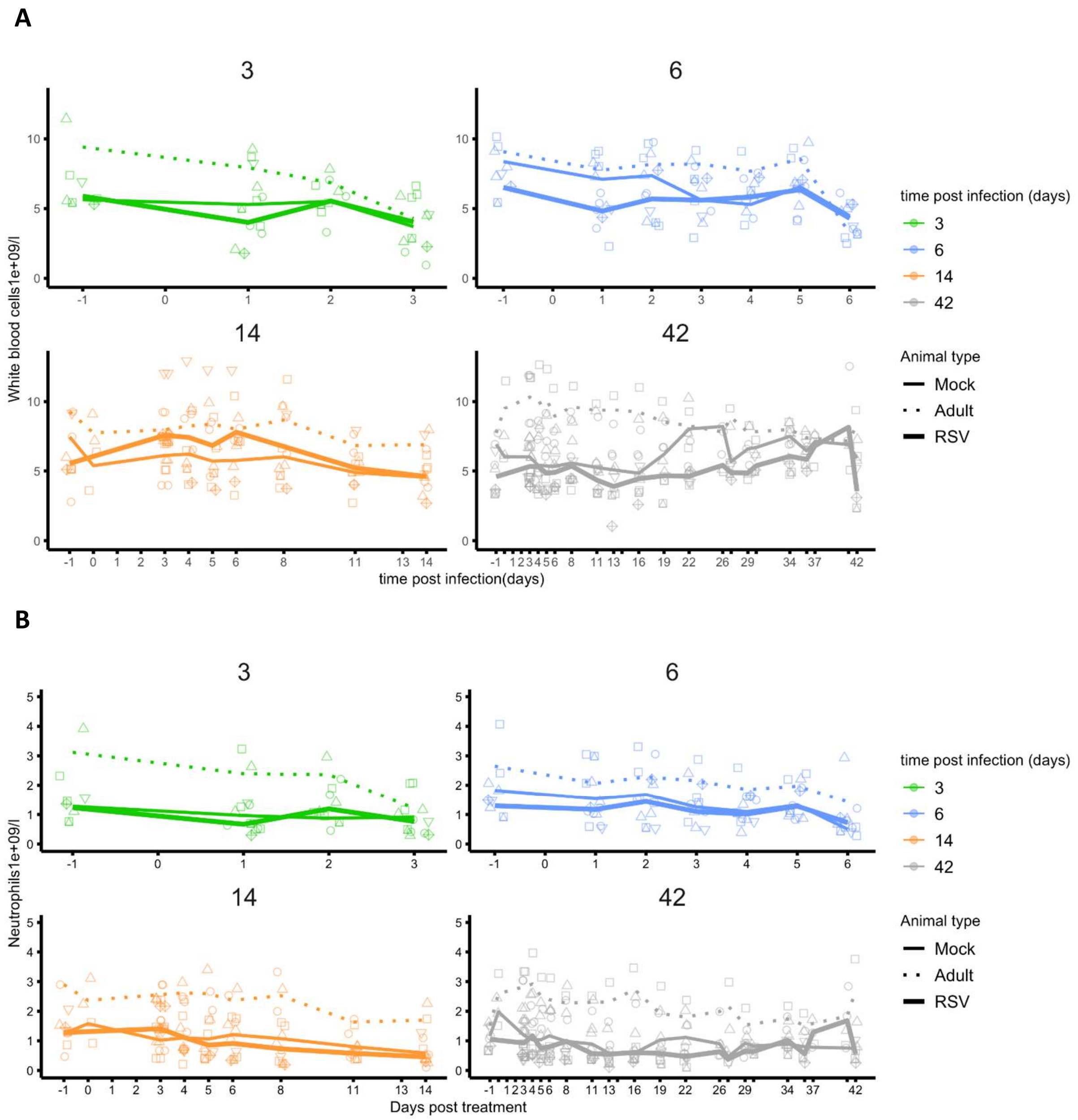


**Figure S1: Longitudinal evaluation of peripheral blood over the course of RSV disease.** Frequencies in the peripheral blood of (**A**) WBC and (**B**) PMNs over the course of RSV disease.

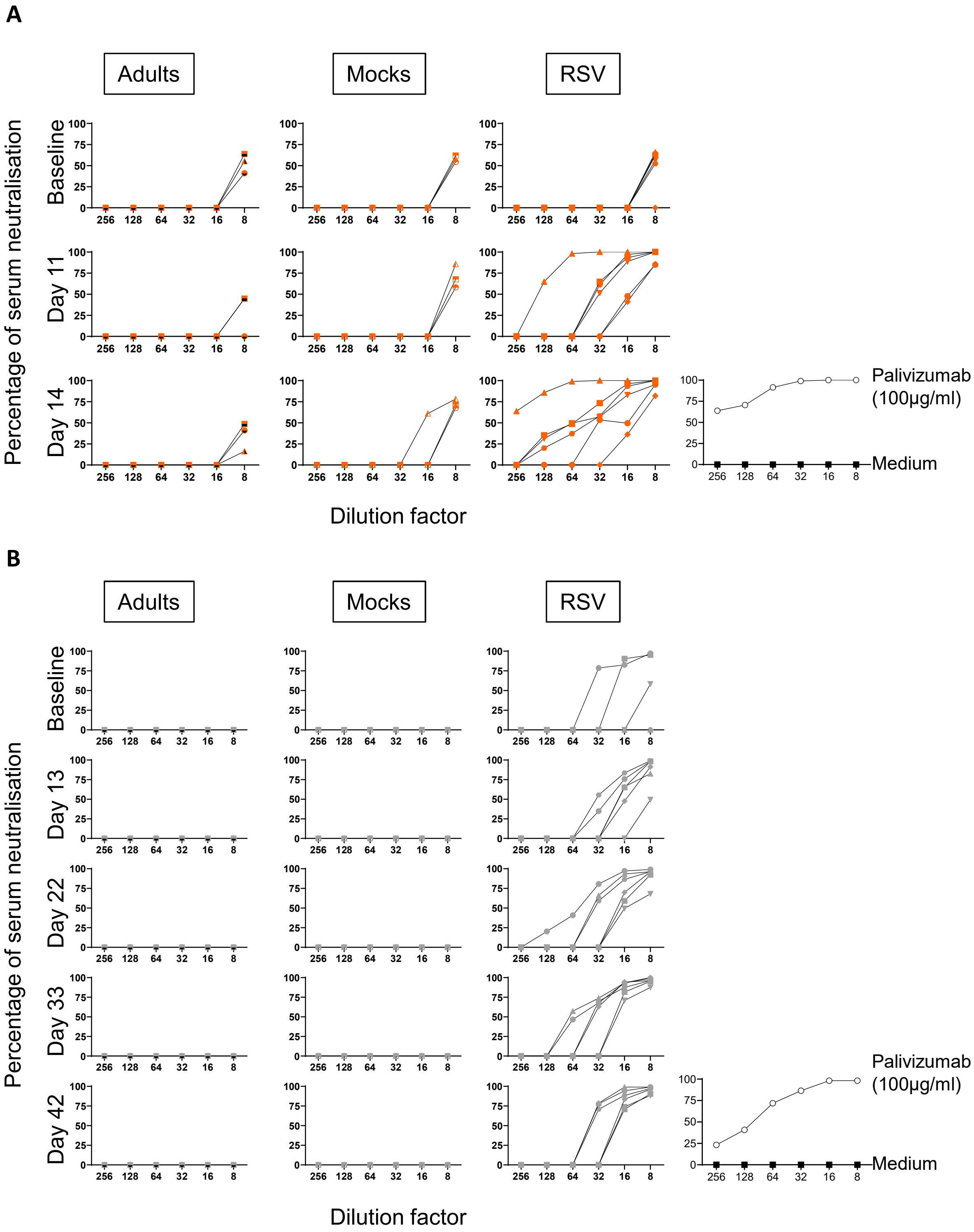


**Figure S2: Evaluation of serum RSV-specific neutralizing antibodies.** RSV-infected neonates have naturally acquired NAb at day 13-14 p.i (**A**), that persist over a 42-day long period (**B**). Each symbol represents an individual animal (symbols filled with black, healthy adults; transparent symbols, mock neonates; solid color symbols, neonates infected with RSV).

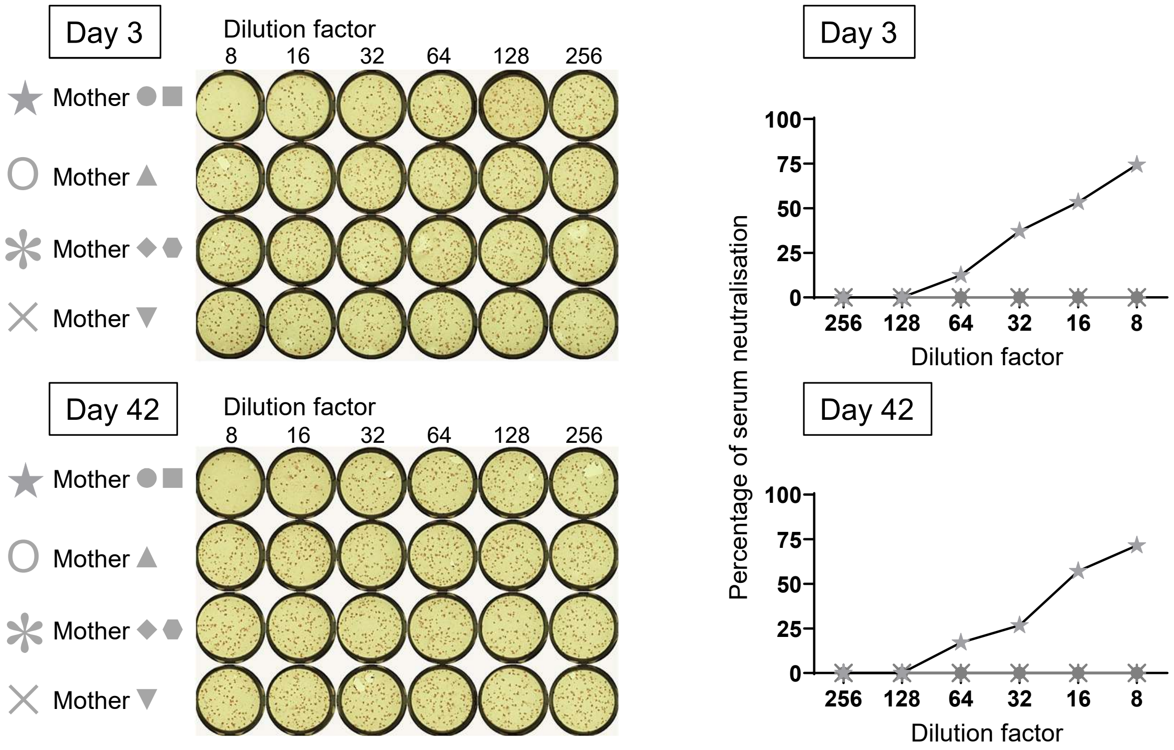


**Figure S3: RSV-specific neutralizing activity present in one uninfected animal.** The mothers of two RSV-infected sibling presented an RSV-neutralization over the 42-day long period of the experiment. Each symbol represents an individual animal.

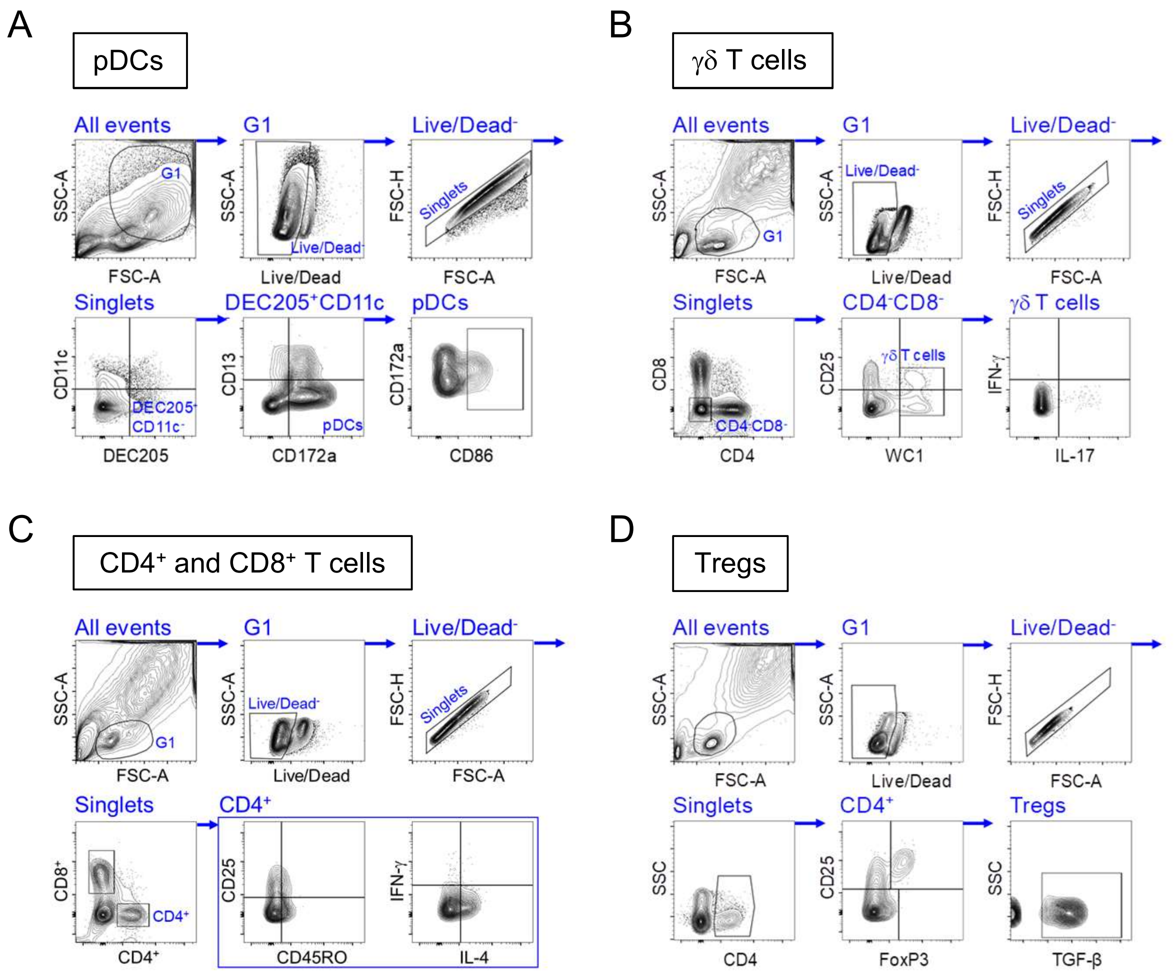


**Figure S4: FCM gating strategy for immune cells identification.** Example of gating strategy for multiparameter FCM analysis of ovine pDCs (A), γδ T cells (B), CD4+ and CD8+ T cells (C) and Tregs (D).

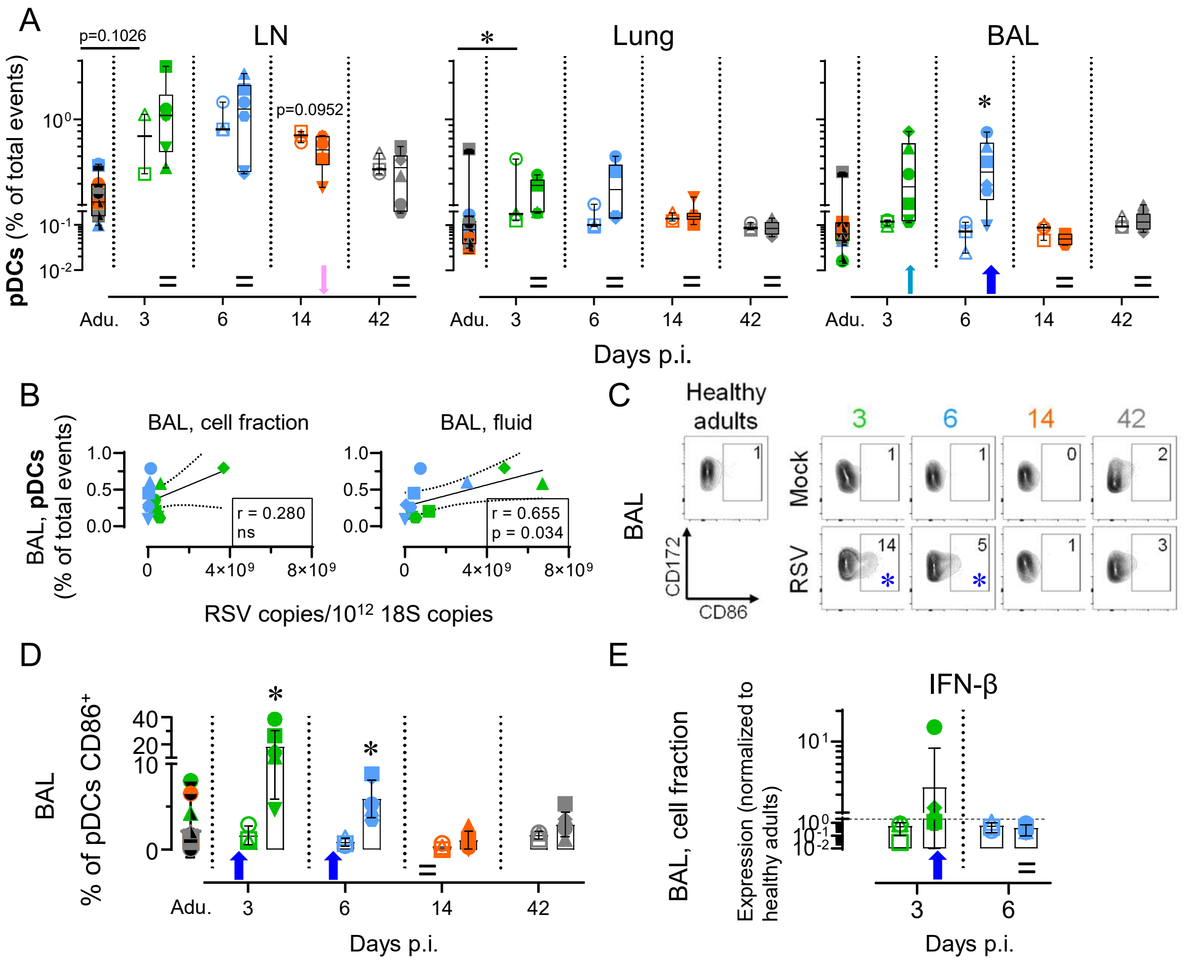


**Figure S5: Neonatal RSV infection induces bronchoalveolar space recruitment of activated pDCs.** (A) Neonatal RSV infection induces an early and transient bronchoalveolar space colonization by pDCs, but not in peribronchial LN or lung tissue. (B) Correlation coefficient (r) obtained with pDC counts calculated as a function of RSV copies/1012 18S copies. Significance was reached for BAL fluid but not for BAL, cell fraction. (C-D) A high frequency of recruited pDCs in bronchoalveolar space upregulate CD86 (D, representative FCM contour plots; E, plot displaying all individuals). (E) Type I IFN quantification by qPCR in the BALs of RSV infected neonates and healthy controls. Each symbol represents an individual animal (healthy adults, n=12; mock neonates, n=3 per time point; neonates infected with RSV, n=6 per time point). Boxplots indicate median value (center line) and interquartile ranges (box edges), with whiskers extending to the lowest and the highest values. Stars indicate significance levels. *, $p < 0.05$.
